# Environmental DNA as a management tool for tracking artificial waterhole use in savanna ecosystems

**DOI:** 10.1101/2020.11.03.367417

**Authors:** Maxwell J. Farrell, Danny Govender, Mehrdad Hajibabaei, Michelle van der Bank, T. Jonathan Davies

## Abstract

Game parks are the last preserve of many large mammals, and in savanna ecosystems, management of surface waters poses a conservation challenge. In arid and semi-arid regions, water can be a scarce resource during dry seasons and drought. Artificial waterholes are common in parks and reserves across Africa, but can alter mammal community composition by favoring drought intolerant species, with consequences for disease dynamics, and population viability of drought-tolerant species. Analysis of waterborne environmental DNA (eDNA) is increasingly used to inform conservation of rare and invasive species, and conduct large-scale biodiversity assessments. To explore the reliability of eDNA as an indicator of mammal waterhole use in savannas, we compare eDNA metabarcoding and camera traps for documenting artificial waterhole use in the Kruger National Park, South Africa, a global hotspot for mammal diversity. We show that eDNA metabarcoding can recover the majority of mammal species detected by camera traps, including a number of endangered species, but DNA signatures of mammal visitation are temporally limited, with best performance when tracking water-dependent large bodied mammals visiting within two days of sampling. Our results highlight limitation of eDNA based monitoring in these systems, including the lack of long-term eDNA persistence in small and highly utilized waterholes, and variability in detection rates among species. However, we demonstrate that eDNA-based approaches can be used to track mammals of conservation concern, and reflect patterns of recent waterhole use and co-occurrence across water-dependent species, both of which are crucial for making evidence-based decisions regarding water management and provisioning.

## Introduction

Across Sub-Saharan Africa, artificial watering holes have been introduced in conservations areas and private reserves (Weir, 1971; Hitchcock, 1996; Berry et al., 2001; Egeru et al., 2015; Chamaillé-Jammes et al., 2016)). These waterholes were intended to increase game numbers by stabilizing water availability year-round and are frequently visited by a diversity of birds and mammals. However, shifting availability of water resources can modify predator-prey interactions (Valeix et al., 2009; Amoroso et al., 2020), influence cross-species pathogen transmission (Turner et al., 2016; Franz et al., 2018), and drive large-scale shifts in community composition (Redfern et al., 2005; Smit et al., 2007). The retention of these artificial sites remains contentious as while they provide opportunities for game viewing that boost important tourism revenue, this can conflict with conservation goals to promote biodiversity and vital ecosystem processes through management strategies that maintain natural spatial and temporal variability in water availability (Smit et al., 2020, 2007).

The Kruger National Park (KNP) is among the oldest game reserves in southern Africa, and a key conservation site for a number of threatened species such as white rhinoceros (*Ceratotherium simum*) (Ferreira et al., 2017), a species of global conservation concern. Poaching and infectious diseases pose severe threats for many species in the park including critically endangered black rhinoceros (*Diceros bicornis*) (Ferreira et al., 2018), hippopotamus (*Hippopotamus amphibius*), elephant (*Loxodonta africana*), and giraffe (*Giraffa camelopardalis*)(Bengis and Erasmus, 1988). The KNP began the introduction of artificial waterholes in the 1930’s, intending to increase game numbers by stabilizing water availability year-round (Redfern et al., 2005). However, these artificial water sources have resulted in the expansion and overabundance of drought-intolerant species such as impala (*Aepyceros melampus*), elephants, and zebra (*Equus burchellii*). Conceived in the era of big-game hunting, the resulting community imbalance led to increased competition for food, negatively impacting rarer drought-tolerant species, such as the sable (*Hippotragus niger*) and roan antelopes (*Hippotragus equinus*), which have suffered marked declines in the park (Smit et al., 2007). Consequently, many artificial waterholes were decommissioned, reducing numbers from over 300 at the peak in the early 1990s (Smit et al., 2007) to roughly 160 in 2015, but their current usage by wildlife and impact on wildlife community structure has not been systematically documented. Successful management of surface waters can be strengthened by applying new tools to conduct biodiversity surveys at waterholes. Here we explore the utility and limitations of two emerging approaches: sequencing of environmental DNA (eDNA), and camera trapping.

Sequencing of eDNA from waterholes and ponds has been used successfully for surveying terrestrial mammals in sub-Saharan Africa (Seeber et al., 2019) and Japan (Ushio et al., 2017), multiple vertebrates in Australian arid zones (Furlan et al., 2020), and both semi-aquatic and terrestrial mammals in the United Kingdom (Harper et al., 2019; Sales et al., 2020b). These studies indicate broad utility, but show that efficacy varies across environmental conditions and species due to differences in abundance, body size, trophic position, drinking behaviour, and tolerance to drought (Ushio et al., 2017; Seeber et al., 2019; Harrison et al., 2019; Harper et al., 2019; Sales et al., 2020a,b; Leempoel et al., 2020; Jo et al., 2020; Lyet et al., 2021). Therefore, despite the promise of eDNA, there remain questions about and how eDNA biomonitoring compares to other survey methods including camera trapping (Harper et al., 2019; Wearn and Glover-Kapfer, 2019; Sales et al., 2020a,b; Leempoel et al., 2020; Lyet et al., 2021).

In contrast to eDNA approaches, camera traps have a longer history of use and have gained increased popularity due to improved picture quality, decreased costs of digital cameras, and progress in citizen science and machine learning for species identification (Norouzzadeh et al., 2018). Camera trap surveys are now also being conducted at larger scales, with savanna ecosystems being the focus of one of the most intensive co-ordinated global camera trap monitoring efforts (Swanson et al., 2015). Camera traps can provide a range of important data relevant for waterhole analyses including abundances, activity patterns, and identification of individuals (Burton et al., 2015). However, camera traps are also subject to numerous logistic challenges including technical failure, theft and animal interference, inconsistent performance across habitats and camera models, and informatic challenges of data storage, curation, and image classification (Glover-Kapfer et al., 2019; Wearn and Glover-Kapfer, 2019). Thus, both camera trapping and eDNA-based approaches have strengths and weaknesses (Harper et al., 2019; Leempoel et al., 2020; Sales et al., 2020a).

Here we evaluate the efficacy of eDNA metabarcoding in documenting waterhole visitation and tracking endangered mammals in the KNP. With support from South African National Parks, we pair camera trap and eDNA approaches to describe the mammal communities associated with artificial waterholes. We contrast camera trap data on species visitation with Cytochrome C Oxidase I (COI) sequences amplified from eDNA present in waterholes, and identify factors influencing the efficacy of eDNA-based approaches. Our study not only explores the utility of eDNA metabarcoding for biodiversity monitoring, but also for quantifying patterns of species co-occurrence, and tracking of endangered species in savanna ecosystems. This work will help inform conservation planning and the management of artificial waterholes and surface water availability in arid and semi-arid regions by evaluating complimentary survey methods, and their utility for future long-term monitoring programs. We suggest eDNA and camera trapping together are likely to become fundamental tools in biodiversity surveys as they can be conducted at greater spatial scales, broader taxonomic scales, and finer temporal scales than previous approaches to document the use of surface waters in these

ecosystems.

## Materials & Methods

### Study Site

Sampling was conducted in the KNP through June and July 2015, when natural sources of surface water are largely dry and waterhole use is highest. Notably, 2015 coincided with an exceptional drought in the KNP and a significant die-off of mega-herbivores (Malherbe et al., 2020). Across the southern half of the park, six concrete bottom artificial waterholes were selected based on distance to the laboratory that allowed for sample processing within 12 hours of collection. These small waterholes hold approximately 2,000L liters of water. Each is equipped with a ball-valve that regulates water levels (Fig. 1). Waterholes varied in shape, with some mimicking the contours of natural pans. Four of the sites were filled with groundwater via boreholes, while two were fed by a pipeline that diverts water from the nearby Olifants river.

**Figure 1:**
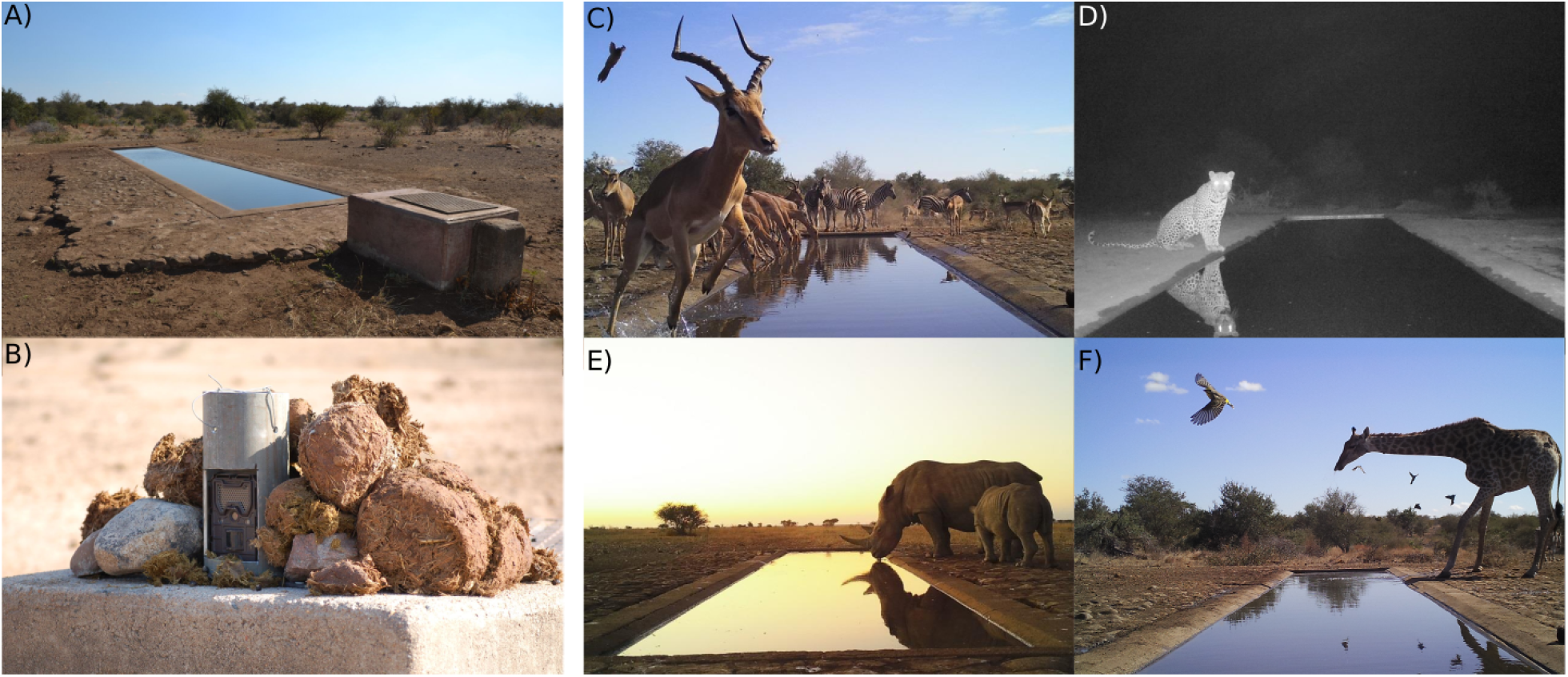
Example of a trough-shaped artificial waterhole (A) with associated camera trap housing attached to the valve housing (B), and examples of photographs (C-F).

### Camera Trapping & Annotation

At each site, 2015 model Bushnell Aggressor Trophy Cam HD Low-Glow cameras were set up either in trees, on poles, or placed in metal cases and attached to the concrete ball-valve housing. Cameras were set to take time lapse photos at five-minute intervals, and also when triggered by motion with a minimum one minute interval between photographs. Individuals present in photographs were annotated to species level, and individuals in contact with the waterhole, or directly adjacent to the waterhole were counted as using the waterhole (animals in the background, or passing through the field of view but without orientation towards the waterhole were excluded).

### Water Sampling

We took water samples after one week and two weeks following camera placement, providing two water samples and two weeks of camera samples per site. At each sampling, we collected two replicate 1L water samples in autoclaved, UV sterilized glass jars from opposite ends of the waterhole. Water samples were placed on icepacks and kept between 4-8° Celsius until until filtering. Because eDNA degradation can be influenced by physicochemical water properties (Jo et al., 2020; Curtis et al., 2021), we also collected data on water temperature (C°), conductivity (mS/cm), dissolved oxygen (% saturation), and pH, using a YSI 650QS multi-parameter sonde.

Collection jars were washed with ELIMINase® (Decon Labs) and rinsed with deionized (DI) water to minimize sample cross-contamination. Because of water turbidity at some sites, filtering the entire collected volume was not feasible. Therefore, from each initial 1L sample, we sub-sampled 150mL for all downstream analyses and filtered this through gamma-irradiated 0.2 *µ*m Supor® hydrophilic polyethersulfone membranes (Pall no. 66234) using 300-mL Advantec polysulfone 47-mm filter funnels and a Pall manifold with vacuum pressure maintained by a Pall filtration pump (model 13158). Prior to filtration, all funnel components and tweezers were sterilized by soaking with 10% bleach for 10 minutes, rinsing with DI water, washing with ELIMINase®, rinsing with DI water, and exposure to UV radiation for a minimum of 30 minutes. To identify potential contamination, negative controls were generated twice during sampling by filtering 1L of deionized water used in the laboratory and subjected to extraction, amplification, and sequencing as for all other samples. Filters were stored in sterile 15mL Falcon tubes and placed in a freezer at -60°C. All samples were filtered within 12 hours of collection.

### DNA Extraction, Amplification, and Sequencing

DNA was isolated from filter papers using MO BIO PowerWater® DNA isolation kits. Three cocktails of primers targeting mammal and vertebrate species (Supplementary Materials (SM) 1.2) tagged with Illumina adapter sequences were used to amplify variable regions of the COI barcode gene – currently the gene with the largest taxonomic coverage across animals (Porter and Hajibabaei, 2018a) – through polymerase chain reaction (PCR). The mam ckt F + R and mam ckt F + 230R primer sets included three new minibarcode primers based on methods described in Meusnier et al. (2008) and modified to better target mammalian species. Primer3 software was used to ensure proper binding properties of the novel primers. The third primer set (BR5) combined forward and reverse primers developed by Hajibabaei et al. (2012) and Gibson et al. (2014).

Two-step PCRs were performed, with Illumina-tagged primers being used in the second step. For each primer set/cocktail, individual primers were pooled (outlined in SM 1.2), and a separate PCR was conducted. A mask was worn during all PCRs, and one reaction was done for each primer set/cocktail (i.e. no replicate PCRs). The primer sets were pooled prior to sequencing. PCRs used a standard mix of 17.8*µ*L molecular grade water, 2.5*µ*L 10× reaction buffer (200mM Tris HCl, 500mM KCl, pH 8.4), 1*µ*L MgCl_2_ (50mM), 0.5*µ*L dNTP (10mM), 0.5*µ*L forward primer (10mM), 0.5*µ*L reverse primer (10mM), 0.2*µ*L Platinum Taq DNA polymerase (Invitrogen), and 2*µ*L DNA as template for a total volume of 25*µ*L. PCRs cycler conditions varied by primer set (see SM 1.2 for details).

Amplification success was confirmed through gel electrophoresis, using a 1.5% agarose gel. PCR products were purified using MinElute PCR purification kit (Qiagen), and quantified through flurometry using a Quant-iT PicoGreen dsDNA assay kit (Invitrogen). Samples were normalized, then multiplexed with the Nextera XT Index kit (96 indexes) (Illumina) and sequenced on an Illumina MiSeq flowcell using a V2 sequencing chemistry kit (2 × 250). In addition the two negative controls of deionized water described above, one negative extraction control, and one negative PCR control were included. The first two negative controls were sequenced, but the extraction and PCR controls were clean and thus sequencing was not necessary.

### Bioinformatic Analyses

Across all samples, we generated a total of 2,709,713 Illumina reads. Sequences were separated by primer set and primer sequences removed using the trim.seqs function in mothur (Schloss et al., 2009). Reads were then processed in R (version 3.5.2) (R Development Core Team, 2008) using the package *dada2* version X (Callahan et al., 2016) following the DADA2 Bioconductor workflow (Callahan et al., 2017b) and the workflow for Big Data (benjjneb.github.io/dada2/tutorial.html). Separate pipelines were developed for each primer set. All reads were filtered by quality, removing sequences with maximum expected error greater than 4 for both forward and reverse reads, and reads with any base pair having Q of 6 or lower. Reads were truncated based on dropoffs in quality profiles. For primer set BR5, reads were truncated to a length of 200 and 150 bp for forward and reverse reads respectively, 230 bp and 140 bp for the mam ckt F + R primer set, 220 bp and 140 bp for the mam ckt F + 230R primer set. As samples were sequenced across four different runs, learning error rates, dereplication, denoising and Amplicon Sequence Variant (ASV) calling (Callahan et al., 2017a) using pooled samples, and merging of paired reads were performed separately for each run. For mam ckt F + R and mam ckt F + 230R merging was done via concatenation. Tables of ASV sequences per sample within each run were then combined and chimera detection using all pooled samples was performed. In total 921,086 reads were retained, representing 2,986 ASVs.

Taxonomic assignment from Phylum to Species was performed using the RDP classifier (Wang et al., 2007) via the *dada2* assignTaxonomy function and a custom KNP Vertebrate COI Reference Library (see SM 1.1 for details) with minimum bootstrap values of 50, 80, 95, and 98. To evaluate the sensitivity of taxonomic assignment to choice of reference library, we also assigned taxonomy using the Porter COI reference library (Porter and Hajibabaei, 2018b), and the MIDORI COI reference library (Machida et al., 2017), both with a minimum bootstrap value of 80%, see Wang et al. (2007).

### Modeling eDNA Detection

We used a hierarchical Bayesian regression with Bernoulli response to model species detections by eDNA metabarcoding across samples. As predictors, we calculated total visitation per species per sample (the number of individuals identified at the waterhole) and the time between sampling and the last instance each species visited a site based on the camera trap photographs. In addition, we included average female body mass as reported in Jones et al. (2009), and environmental predictors including temperature, conductivity, dissolved oxygen, and pH. We also included hierarchical predictors for site and species effects, each with adaptively regularizing priors to allow for partial pooling across groups (see SM 1.3 for model formula).

We fit the model in Stan 2.18.0 (Carpenter et al., 2017) via the R package *brms* 2.7.0 (Bürkner, 2017). We assumed weakly informative priors assessed through prior predictive simulation. The model was run across four chains, with 10,000 iterations per chain and the first 5,000 iterations discarded as burn-in and remaining iterations thinned to retain every fourth iteration. Convergenge was checked by visual inspection of traceplots and Rhat equal to 1.0 for all estimated parameters. Model fit was assessed using posterior predictive checks.

### Species Co-occurrence Patterns

To explore patterns in species co-occurrence, we investigated overlap in waterhole use as determined by eDNA metabarcoding and camera trapping. For eDNA, we counted the number of samples in which each species pair was detected. To match this with camera trap data, we counted the number of site-weeks in which pairs of species were detected with camera trapping, throughout the sampling period, and subset to 36 hours before water sampling (see Results). Our approach is not an attempt to quantify the physical co-occurrence of species pairs at an exact time and location, but rather identifies a temporal window in which pairs of species overlap in space.

## Results

### Taxonomic assignment

Total detected species richness from eDNA ranged between 13 and 30 species across reference libraries and RDP minimum bootstrap values (Fig. SM3, Table SM2). The previously compiled COI reference libraries for chordata (Porter–”TP”) and metazoans (MIDORI–”MID”) identified fewer species compared to our custom library (Fig. SM3, Table SM2). Taxonomic assignment using our curated library and a minimum bootstrap of 50% (KNP50) returned 30 species. These largely comprised mammals (19 species), but also included seven birds, two reptiles, one amphibian, and one fish species (Fig. 2, Table SM2). Increasing the minimum bootstrap value to 80% returned fewer species (14 mammals and two birds), with no species-level detections for reptiles, amphibians, or fish. For all subsequent analyses we used the KNP vertebrate reference library with an 80% minimum bootstrap (KNP80) as this provided the same species richness as higher bootstrap values using the same reference library (Fig. SM3), but only included mammals that were documented in the camera traps, thus minimizing false positives – errors of commission. Using the KNP80 approach, negative controls identified a total 54 ASVs (5,066 reads total) across both controls, eight of which were present in both negative controls. However, 52 ASVs could only be assigned to Phylum level, while the remaining two could only be assigned to Class. Since no ASVs found in the negative controls could be assigned to species level, we concluded no significant cross-contamination, and thus there was no need to use the negative controls to filter ASVs identified in the rest of our samples (as all of our subsequent analyses involve analysis of sequences identified to the species level).

**Figure 2:**
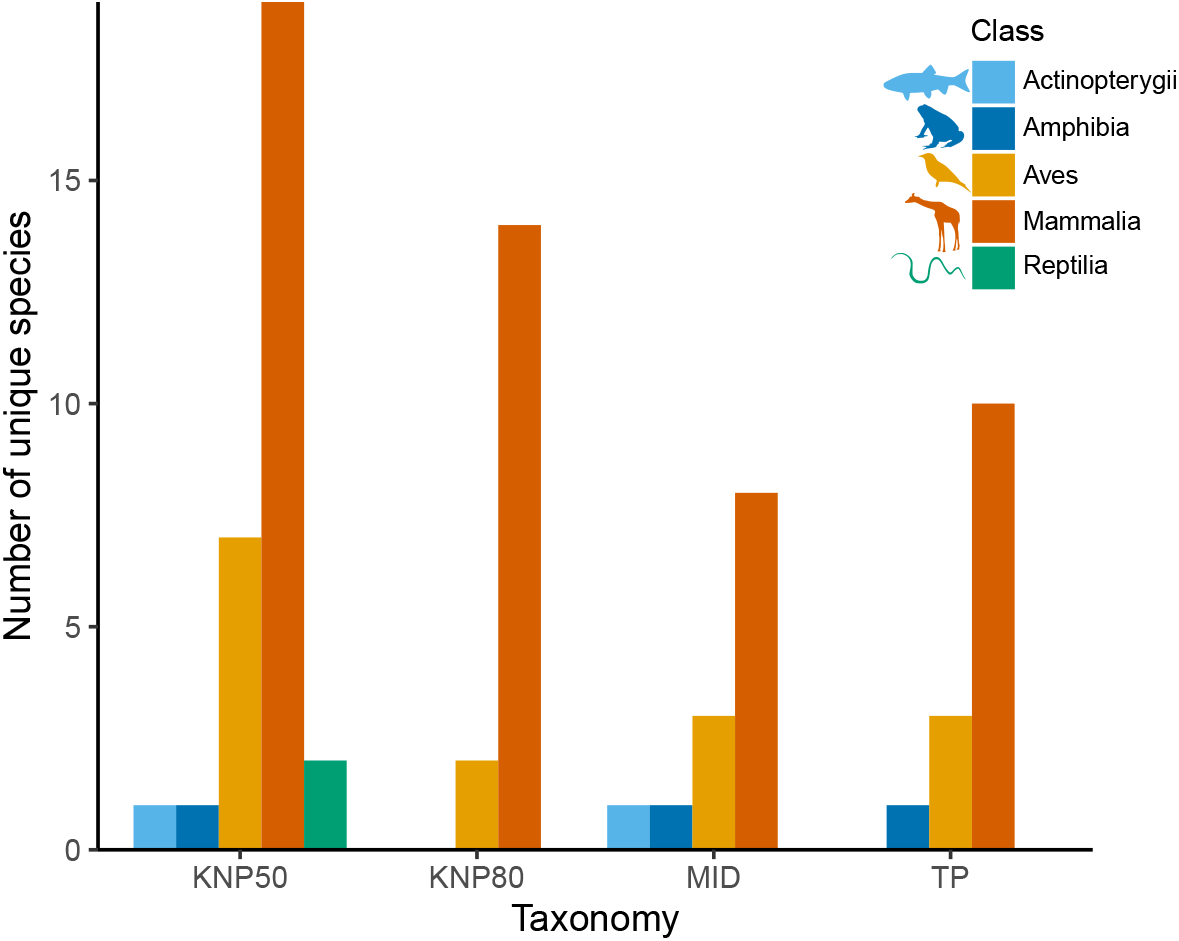
Number of species detected grouped by Class, according to the KNP reference library at the 50 (KNP50) and 80 % minimum bootstrap values, and the Porter Chordata reference library (TP) and the MIDORI reference library (MID) at 80% minimum bootstrap values. The MIDORI and Porter reference libraries returned single fish and amphibian species which were different than those identified by the KNP50 approach (Table SM2) and not known to be present in the KNP.

### Comparing eDNA and camera trap detections

We collected 11 weeks of uninterrupted camera trapping with associated water samples, resulting in 16,027 annotated photographs. We identified 21 mammal species by camera trap, 14 of which were detected by eDNA (Fig. 3). Two species documented by camera traps (*Mungos mungo* and *Paraxerus cepapi*) were not detected by eDNA because they did not have representative sequences in the KNP vertebrate reference library. Across all species with reference sequences, 25% of species by sample combinations identified by camera were recovered by eDNA. When restricting to only species seen in the last 12 hours before sampling, 50% of species by sample combinations identified by camera were detected by eDNA. When comparing shorter camera trap sampling times (3-12 hours before eDNA sampling), cameras and eDNA return similar diversity estimates (Fig. SM1). While we observed a negative relationship between total visitation and time since last visit of a species, species last visiting a site *>*36 hours before sampling were rarely detected, irrespective of overall visitation frequency (Fig. 4).

**Figure 3:**
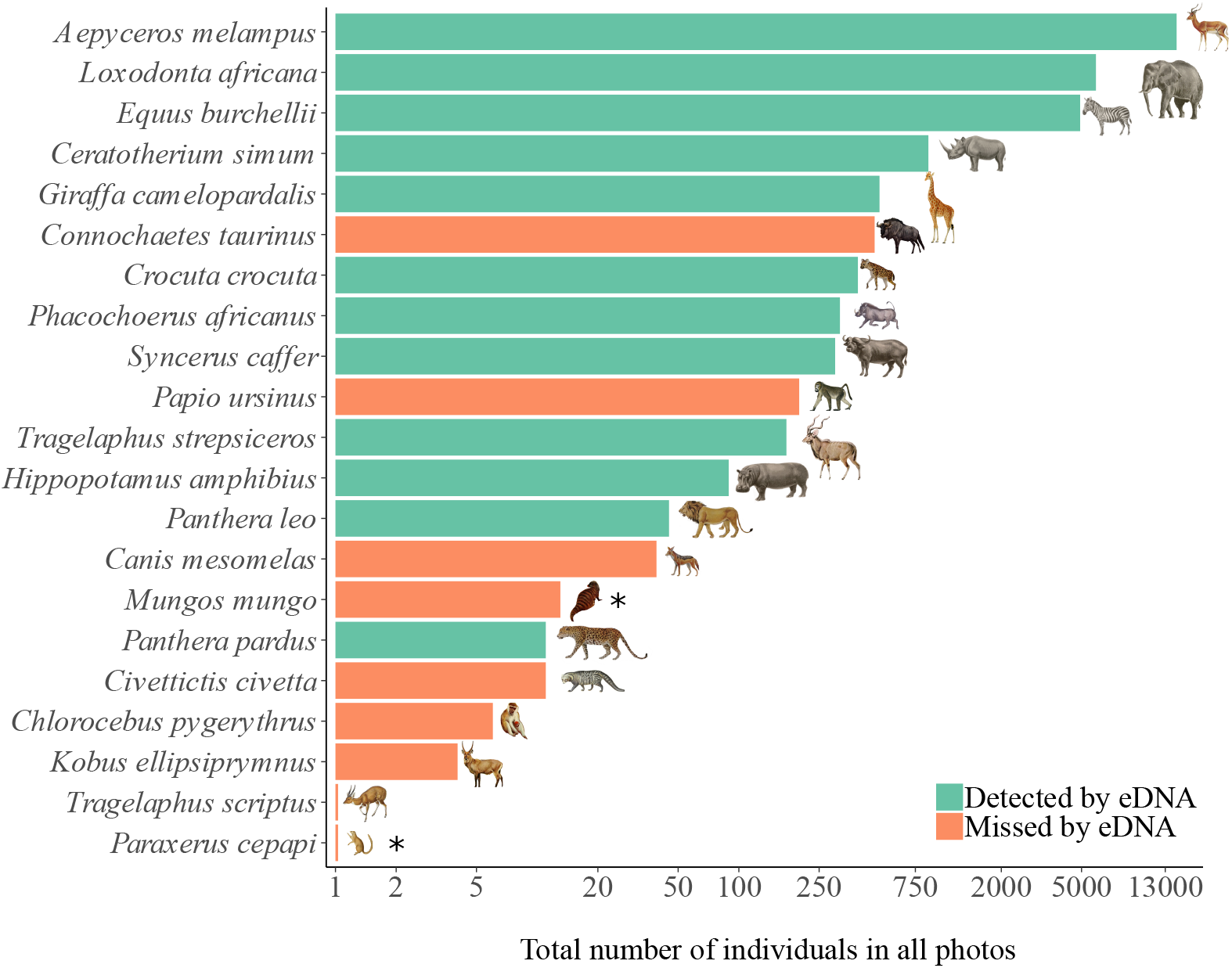
Number of individuals per species across all photographs. Coloring represents detection by eDNA across any of the samples. *species without representative sequences in the KNP vertebrate reference library.

**Figure 4:**
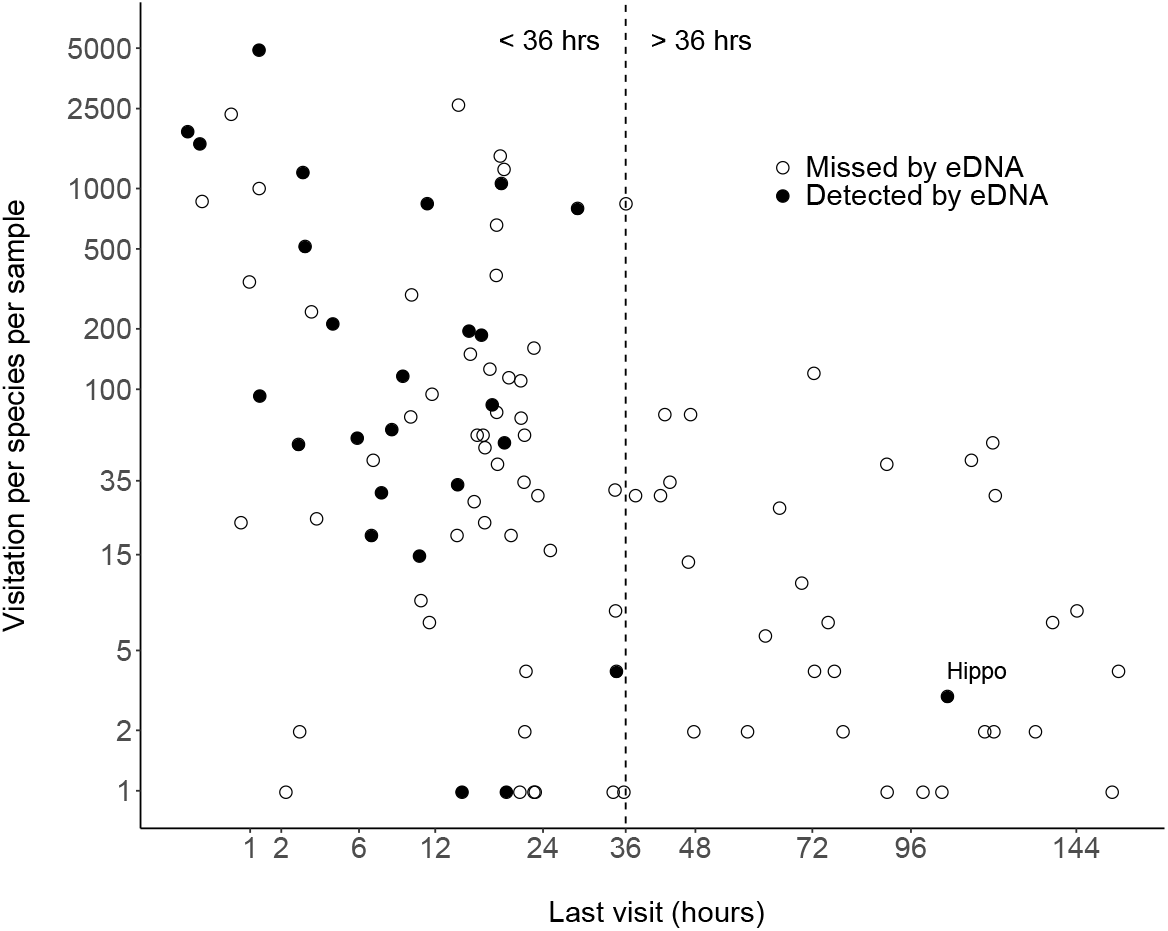
Raw data plot showing the relationship between the number of hours before sampling that a species last visited the site and the total count of visitations per sample over the previous week. Filled points indicate samples which have positive eDNA detection for the given species.

### Variation in eDNA detection among species

We found large variation in the raw eDNA detection rates among species (Table SM4). Two species (baboons (*Papio ursinus*) and wildebeest (*Connochaetes taurinus*)) were never detected by eDNA despite visiting waterholes in relatively high numbers (Fig. 3). The clearest species-level predictor of detection probability was time since last visit (Fig. 5A-B, Table SM3), and there was a rapid drop in detection probability after 36 hours (Fig. 4). Body mass was also an important predictor of detection success. While smaller bodied species visit waterholes less often than larger mammals (Fig. SM4), body mass also had a relatively large positive effect in our eDNA detection model adjusting for total visitation and time since last visit, (Fig. 5A, Table SM3), indicating that larger species may have behaviours or physiologies that make them shed eDNA at higher rates. We were never able to detect eDNA from species with an average mass less than 50 kg.

**Figure 5:**
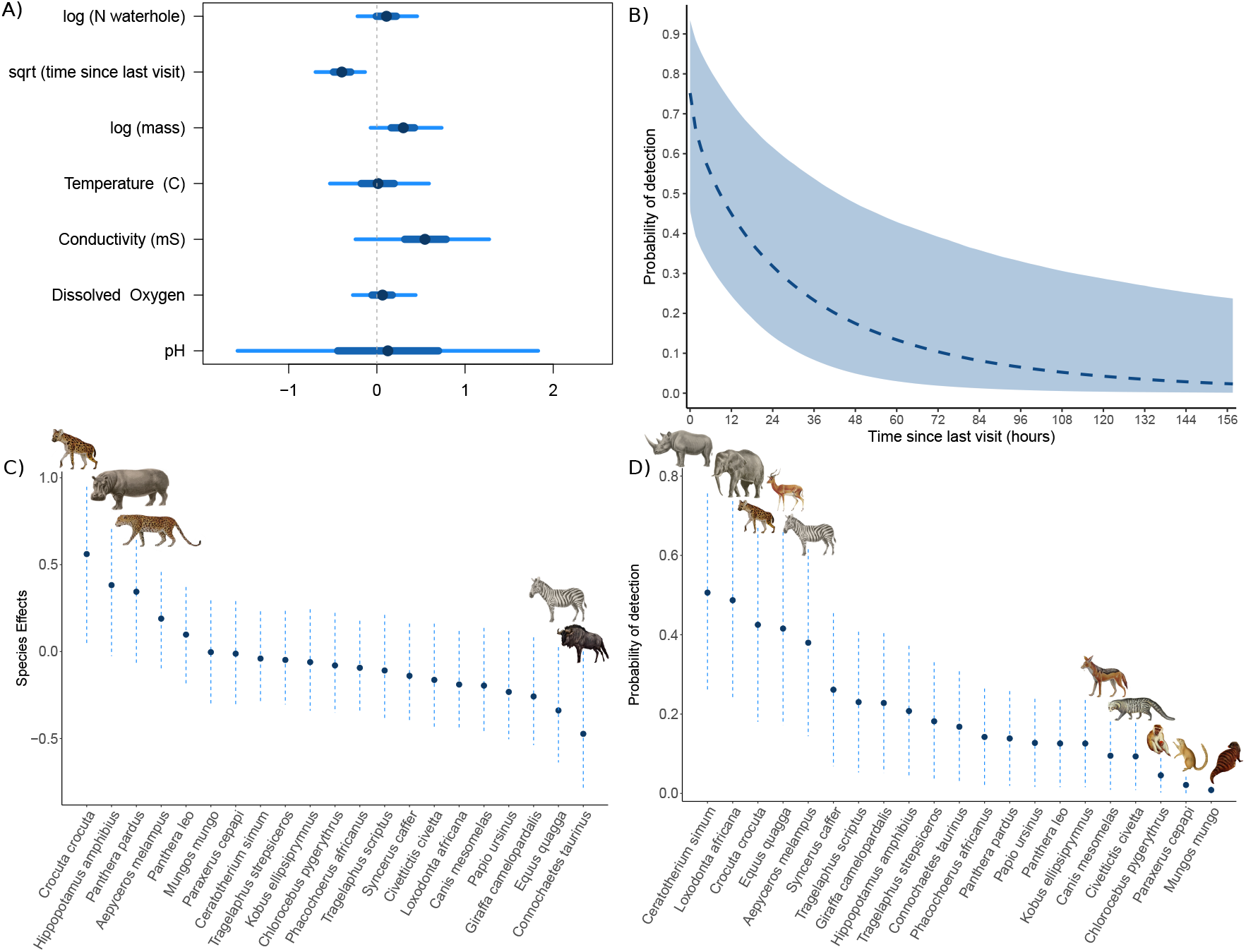
A) Estimated regression coefficients for continuous predictors from the eDNA detection model. Points represent mean estimates; thick and thin lines represent 50% and 95% credible intervals respectively. B) Conditional effect of time since a species last visit on the probability of eDNA detection. Conditional effects are estimated using the mean for continuous variables. The dashed line represents the mean effect, with the shaded area indicating the 95% credible interval. C) Estimated hierarchical effects for species, presented on the log-odds scale. These indicate that hyena and hippopotamus are detected more often than we would expect from their size and waterhole visitation patterns alone. D) Posterior predictions of species detection by eDNA metabarcoding – these predictions show that, on average, white rhino and elephant are the species most likely to be detected in any given water sample. For panels C and D, points indicate posterior means, with bars representing +/- the variance and species are arranged by decreasing mean. Animal images are used to help illustrate top and bottom ranked species in these panels.

We used posterior predictions from our model to characterize the mammalian waterhole community as detected by eDNA (Fig. 5D). Unlike the species-level effects, these represent species detection rates based on predictions from the fit model. We find that white rhinos (*Ceratotherium simuim*) and elephants (*Loxodonta africana*) have the highest mean probabilities of being detected, followed by hyena (*Crocuta crocuta*), zebra (*Equus quagga*), and impala (*Aepyceros melampus*). Wildebeest (*Connochaetes taurinus*) and baboons (*Papio anubis*), which were never detected by eDNA, have posterior probabilities of detection comparable to other species, indicating that given their waterhole ecology it is unexpected that we did not find a positive eDNA detection in our study. While baboons fell below the approximate size threshold (50kg) for detection in this system, we were surprised not to detect wildebeest as they are large bodied and appear to have visitation patterns similar to other large antelopes that we detected via eDNA. Primer bias is one of the largest sources of taxonomic bias in eDNA metabarcoding studies (van der Loos and Nijland, 2021; Peixoto et al., 2021). We conducted *post hoc* BLAST matches between our primers and archived COI sequences for wildebeest, which suggested strong alignment, but it is possible other molecular biases, such as sub-optimal annealing temperatures for this species (van der Loos and Nijland, 2021), limited our ability to amplify these sequences.

### Waterhole Activity Patterns

Camera traps showed that species have large differences in the timing of waterhole use across weeks (Fig. SM6), likely reflecting species specific differences in water needs. We also detect this variation in species co-occurrences via our eDNA profiles (Fig. SM7), though the lower species richness and detection rates of eDNA only allow for a coarser understanding of these patterns. Comparing species co-occurrences inferred via eDNA and camera trap data subset to photographs taken less than 36 hours before water sampling (removing bias introduced by the temporal decay in eDNA detection), and for only species detected by both methods, there is a correlation of 0.57 (p*<*0.001) between the two co-occurrence matrices (Fig. 6A-B).

**Figure 6:**
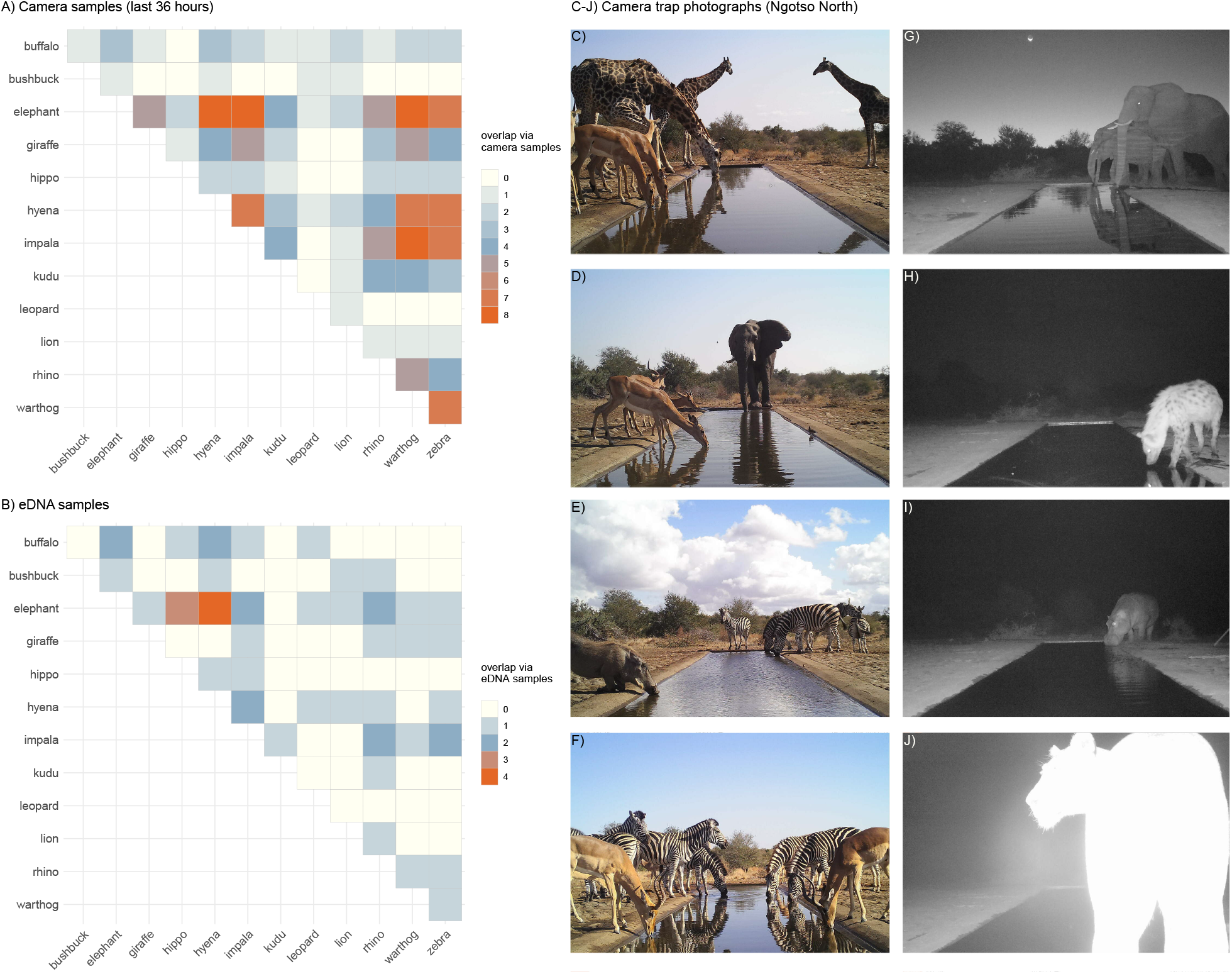
Comparison of overlap in waterhole usage documented by (A) camera traps (restricted to detections in the 36 hours prior to water sampling) and (B) by eDNA metabarcoding. After subsetting, the correlation between the two heatmaps is 0.57 (p*<*0.001). Camera trap photographs from one site (Ngotso North) illustrate species co-occurrences (C-F) and single species visitations which are common during hours of darkness (G-J).

## Discussion

In the KNP, provisioning of water via reservoirs and artificial waterholes are a key management issue (Smit et al., 2007; Redfern et al., 2005). Artificial waterholes have promoted water-dependent species such as impala, zebra, and elephants, largely to the detriment of many drought-tolerant species which have suffered from competition for forage (Harrington et al., 1999). Our camera trapping confirms that waterholes in the KNP are dominated by these three water-dependent species, and demonstrate that eDNA metabarcoding can recover the signature of waterhole visitation for medium to large mammals, accurately capturing the water-dependent mammal community. Our results reinforce the growing consensus that artificial waterholes contribute to the taxonomic dissonance of wildlife communities in arid lands, and we highlight the efficacy of waterhole eDNA for wildlife monitoring. While frequent waterhole use make elephants, impala, and zebra good targets for eDNA detection, the drinking ecology of several other species, including rhinos and hyenas, also make them promising targets for eDNA-based monitoring. However, we find that eDNA performance also varies across species, is sensitive to the choice of genomic reference library and bioinformatic pipeline, and that detection rates can rapidly decay over time.

Compared to eDNA metabarcoding, camera traps offer unparalleled detail on species abundances, temporal visitation patterns, co-occurrences, and the potential for the identification of individuals (Burton et al., 2015). However, we show that eDNA metabarcoding can also provide a useful index of spatio-temporal overlap of species, which may be used to inform management of waterborne diseases in artificial surface waters. In addition, eDNA approaches offer many unique advantages. Water sampling requires a very small amount of time in the field, meaning that multiple distant sites can be covered in a single day. Importantly, eDNA-based approaches also allow for the identification of cryptic species, such as pathogenic microbes (see Farrell et al. (2019)). We suggest the two approaches are thus highly complementary, and when used together can track trajectories of communities over time, providing rich data to help guide applied conservation management decisions in the KNP and other savanna ecosystems across sub-Saharan Africa.

The long history of conservation and biodiversity monitoring in the KNP allowed us to start with complete species lists for vertebrate groups in the park, and generate a curated set of reference sequences. We show improved performance of eDNA detection across species when using our custom reference sequences as compared to existing reference libraries. Our results caution against using pre-configured reference libraries for eDNA-based biodiversity assessments, especially in areas where comprehensive species lists may be obtained. We suggest that future eDNA metabarcoding efforts focus on leveraging expertise of park scientists and existing natural history collections to improve sequence reference libraries. As methods for sequencing DNA from preserved specimens increase in sensitivity (Prosser et al., 2016), local collections present an increasingly valuable DNA source for expert-identified samples.

We found eDNA detection to be temporally limited in our system, with highest detection probability for species visiting waterholes within 24-36 hours of sampling. For example, some species that infrequently visit waterholes, such as lion, are successfully detected when they visit within 24 hours prior to sampling, while other species that visit in higher numbers may be missed by eDNA simply because they did not visit just prior to sampling (see Fig. SM5 for illustrative time series). We observed intense usage of these waterholes, with near complete emptying in some cases, and it appears that previously deposited eDNA is being actively removed through subsequent drinking, rather than molecular degradation. Thus increasing both the frequency of sampling and the volumes filtered would be needed to increase the rate of species detections with eDNA sequencing. In addition, it may be possible to target particular species by sampling at times of day that coincide with daily activity patterns.

Beyond time since last visit, we identified additional species variation in eDNA detection. Hyena, hippopotamus, leopard, impala, and lion were estimated to have slightly elevated species effects, indicating that we have a higher probability of detection. Conversely, we find a lower probability of detecting wildebeest, zebra, giraffe, and baboons. The species level effects capture other qualitative observations, most obviously, our inability to detect wildebeest, and species with relatively small body mass (*<* 50kg), such as baboons, jackals, civets, and vervet monkeys, despite having representative sequences in our reference library. Low detection rates may reflect molecular limitations, such as primer bias, environmental factors such as water conductivity (Collins et al., 2018), and/or unmeasured ecological or behavioural traits, such as the potential for species to be “messy” or “clean” drinkers which influences rates of DNA shedding. Because of such intrinsic site and species differences, it is likely that even well-designed sampling strategies will miss some species, and in systems where there is uncertainty about true species occurrences, absence of evidence should not be taken as evidence of absence.

We attribute the temporal decay of eDNA in our study in large part to the overall visitation rates and rapid removal and turnover of water through drinking. In the Kruger, artificial waterholes are a key management issue as they have led to an overabundance of drought-intolerant species to the detriment of many drought-tolerant (Smit et al., 2007). Through camera trapping we show that the artificial waterholes in the KNP are dominated by three drought-intolerant species (impala, zebra, and elephants), which had orders of magnitude higher visitation rates compared to other species observed at the waterholes. While this extreme skew in waterhole use is likely reducing the ability for eDNA to capture longer term dynamics of waterhole visitation, the drinking ecology of other species, such as rhinos and hyenas, make them good targets for eDNA sampling. Even with the imperfect detection of eDNA metabarcoding, it provides an effective method for sampling a large number of sites, requiring less time in the field, and with less potential for mechanical failure or tampering, as was common with our camera traps.

While high temporal turnover of eDNA in these sites may necessitate increased sampling frequency to capture total species richness at a site, it offers a unique opportunity to study cryptic species associations. We find a significant, if noisy, correlation when comparing detections by both eDNA and camera trapping across the full time window, which increases when restricting to animals seen within 36 hours before sampling, indicating that eDNA may provide a crude proxy for species spatial overlap within time windows relevant for potential cross-species microbial or pathogen transmission via waterholes (see (Farrell et al., 2019)). We suggest that a large proportion of microbial taxa in these waterholes may be animal-associated, representing pathogens, or part of the oral, fecal, or skin microbiomes of visiting host species. In addition to pathogens, artificial waterholes may be sources for horizontal transfer of mutualist or commensal taxa, or antibiotic resistance plasmids carried from outside of the park (Mariano et al., 2009). Thus, while fine-scale visitation patterns are better assessed by camera traps and the short persistence time of eDNA in waterholes limits our ability to characterize waterhole usage beyond a few days of activity, eDNA sequencing offers an opportunity to use non-invasive sampling to identify host-associated microbes, and examine how water provisioning may be impacting pathogen and microbiome dynamics.

Our study illustrates the utility of eDNA as a tool for biodiversity monitoring, and how it offers an added understanding of species ecology through pairing with camera trapping. As methods improve with respect to choice of marker gene, primers, reference libraries, and sequencing depth we expect increased detection ability (Ushio et al., 2017; Lyet et al., 2021), but similar biases to those highlighted here will likely persist. Nonetheless, we have shown that easily-collected water samples can provide reliable data for tracking species of conservation concern in the KNP, such as rhinoceros Ferreira et al. (2015), and the critically endangered white-backed vulture (Murn et al., 2013). For the Kruger and other African savanna ecosystems, we hope our study lays the groundwork for future exciting work exploring cryptic biodiversity, the impacts of water provisioning on rare and threatened taxa, and the dynamic nature of species associations in these vital conservation areas.

## Data & Code Accessibility

Scripts to generate the reference library, format public references for use with DADA2, perform all analyses, and generate the figures are available at github.com/maxfarrell/eDNAcamtrap (archived at 10.5281/zenodo.6977480), along with data necessary to reproduce the analyses. Raw sequence data per sample are available via the Sequence Read Archive (SRA BioProject PRJNA490450).

## Author Contributions

MJF and TJD designed the study with input from all co-authors. MJF gathered the data and performed water filtrations with support from DG and MvdB. MH designed the molecular workflow and performed DNA extractions, amplifications, and sequencing. MJF designed and performed all data analyses. MJF wrote the paper with input from TJD and DG.

## Acknowledgements

This work would not have been possible without logistical support from the African Centre for DNA Barcoding and Purvance Shikwambana, molecular work by Shannon Eagle, photographic identifications by Cody Danaher, species inventories from the Kruger National Park Museum staff, and the knowledge and protection provided by Kruger game guards Thomas, Velly, and Martin. We also thank Steven Kembel, his lab, Alexis Carteron, Joanne Littlefair, and Benjamin Callahan for constructive feedback on bioinformatic analyses. Special thanks to B.S. Bezeng for his friendship, dedication, and long days in the field. This work was conducted with support and permission from SANParks (project DAVIJ1256). MJF was supported by a Vanier NSERC CGS, the Vineberg Fellowship, the CIHR SBTP, and project funding from the Quebec Centre for Biodiversity Science, McGill Biology, and an NSERC Discovery Grant awarded to TJD.

## Supplementary Materials

### 1 Supplementary Methods

#### 1.1 Assembling the Kruger Vertebrate COI Reference Library

We assembled a COI KNP reference library by first compiling complete species lists for mammals, birds, reptiles, amphibians, and fish maintained by the Kruger National Park Museum, returning a total of 854 named species. We then identified species synonyms via the *taxize* package in R (Scott Chamberlain and Eduard Szocs, 2013; Chamberlain et al., 2020) returning 1,283 Latin binomials. We downloaded representative COI sequences from GenBank via the R package *rentrez* using the following search string in the nuccore databate: “COX1[gene] OR cox1[gene] OR coxI[gene] OR CO1[gene] OR COI[gene] OR Cytochrome c oxidase subunit I[gene] OR cytochrome c oxidase subunit I[gene] OR cytochrome oxidase subunit 1[gene] OR Cytochrome oxidase subunit 1[gene]) AND 0:5000[Sequence Length] AND *SPECIES* [ORGN]”, where *SPECIES* was replaced with the species Latin binomial. Since barcode sequences available in the BOLD public database (Ratnasingham and Hebert, 2007) are not always found in GenBank (Porter and Hajibabaei, 2018), COI-5P sequences were also downloaded from BOLD via the R package *bold* (Chamberlain, 2017). Finally, to gather additional COI sequences for species which were not available via the search strategies described above, we excised COI sequences from whole annotated mitochondrial genomes available on GenBank using the *PrimerMiner* R package (Elbrecht and Leese, 2017) functions *Download mito* and *Mito GB2fasta*. To clean and format the reference sequences for use with *dada2*, we concatenated individual FASTAs per species into a single file, changing multi-line FASTAs to single-line, removing leading or trailing ambiguous bases (denoted by “N” or “-”), and excluding sequences that still included ambiguous base assignments.

We performed manual cleaning and curation of the concatenated FASTA sequence matrix, which involved sequence alignment via MUSCLE (Edgar, 2004) including an additional reference sequence used to identify the Folmer region (GBMA3852-12). After alignment, reference sequences were trimmed to the Folmer region in Aliview (Larsson, 2014), then the Folmer reference sequence and any duplicate sequence removed. Sequences were then realigned with MUSCLE and poorly aligned sequences removed. Alignment gaps were removed and sequences were filtered to include only those longer than 249 bp. We then performed another iteration of removal of duplicated sequences, alignment, manual cleaning of poorly aligned sequences, and removal of alignment gaps. We used this output as our custom KNP vertebrate COI reference library. We next generated a dada2-compatible taxonomy mapping file based on the KNP species inventories. Scripts (bash and R) to download sequences and format the resulting set of FASTAs are available in the Data & Code Supplement (to be archived via Zenodo upon acceptance), and at github.com/maxfarrell/eDNAcamtrap.

##### 1.1.1 Outline of the KNP Reference Library Assembly

This is a textual outline for generating the Kruger Vertebrate COI Reference Library. For scripts to reproduce the automated sections of this workflow see the Data & Code supplement (github.com/maxfarrell/eDNAcamtrap, to be archived via Zenodo upon acceptance).

1. Mammal latin binomials and taxonomy from KNP inventories
2. Identify synonyms with R package *taxize*
3. Download COI sequences from Gen Bank (via R package *rentrez*) *Search term in nuccore database:* “COX1[gene] OR cox1[gene] OR coxI[gene] OR CO1[gene] OR COI[gene] OR Cytochrome c oxidase subunit I[gene] OR cytochrome c oxidase subunit I[gene] OR cytochrome oxidase subunit 1[gene] OR Cytochrome oxidase subunit 1[gene]) AND 0:5000[Sequence Length] AND *SPECIES* [ORGN]” where *SPECIES* was replaced with the species Latin binomial.
4. Download COI sequences from BOLD public database (via R package *bold*) Search for species name in and pull COI-5P sequences.
5. Excise COI sequences whole mitochondrial genomes using (via R package *PrimerMiner* functions Download mito and Mito GB2fasta)
6. Merging and cleaning reference library
  a. Concatenate all individual fastas per species into one file
  b. Change multi-line fasta sequences to single-line
  c. Remove trailing ambiguous bases (N or -)
  d. Remove leading ambiguous bases (N or -)
  e. Remove sequences that have any ambiguous bases (N or -) and the preceding header line
  f. Generate taxonomy mapping file
  g. Prune FASTA sequences to only those in the taxonomy
  h. Remove any duplicated sequence IDs in the fasta (they seem to come from GB)
  i. Remove and lines with only “*>*“
  j. Alignment with Muscle for visual check of sequences
  k. Add reference to identify folmer region (GBMA3852-12)
  l. Alignment with Muscle
  m. Manual trimming of sequences to folmer region in AliView
  n. Remove duplicated sequences
  o. Alignment with Muscle
  p. Manual cleaning of poorly aligned sequences in AliView
  q. Alignment with Muscle
  r. Manual cleaning of poorly aligned sequences in AliView
  s. Remove alignment gaps
  t. Filter sequences to those longer than 249 bp
  u. Alignment with Muscle
  v. Remove duplicated sequences
  w. Alignment with Muscle
  x. Manual cleaning of poorly aligned sequences in AliView
  y. Remove alignment gaps
  z. save as “final_Oct15_2019.fasta”

Our curated reference library contained 4965 unique sequences for 391 species. This library covers 86 of the 147 mammals documented in the park, and although coverage is low for some small-bodied and diverse orders (23/41 Chiroptera, 7/23 Rodentia, 1/9 Insectivora), the orders most likely to actively use the waterholes are well represented (23/26 Cetartiodactyla, 17/27 Carnivora, all 5 Primates, and all 3 Perissodactyla) (see Table SM1 for counts by mammalian order).

**Table SM1:** Counts of mammal species per order included in the Kruger Vertebrate Reference Library, present in the park according to KNP Library species lists, and the percentage of species in the park which are represented in the reference library.

| Order | Species in library | Species in park | % Represented |
| --- | --- | --- | --- |
| *Carnivora* | 17 | 27 | 63 % |
| *Cetartiodactyla* | 23 | 26 | 89 % |
| *Chiroptera* | 23 | 41 | 56 % |
| *Hydracoidea* | 1 | 2 | 50 % |
| *Insectivora* | 1 | 9 | 11 % |
| *Lagomorpha* | 1 | 3 | 33 % |
| *Macroscelidea* | 2 | 3 | 66 % |
| *Perissodactyla* | 3 | 3 | 100 % |
| *Pholidota* | 1 | 1 | 100 % |
| *Primates* | 5 | 5 | 100 % |
| *Proboscoidea* | 1 | 1 | 100 % |
| *Rodentia* | 7 | 23 | 30 % |
| *Tubulidentata* | 1 | 1 | 100 % |

#### 1.2 Primers

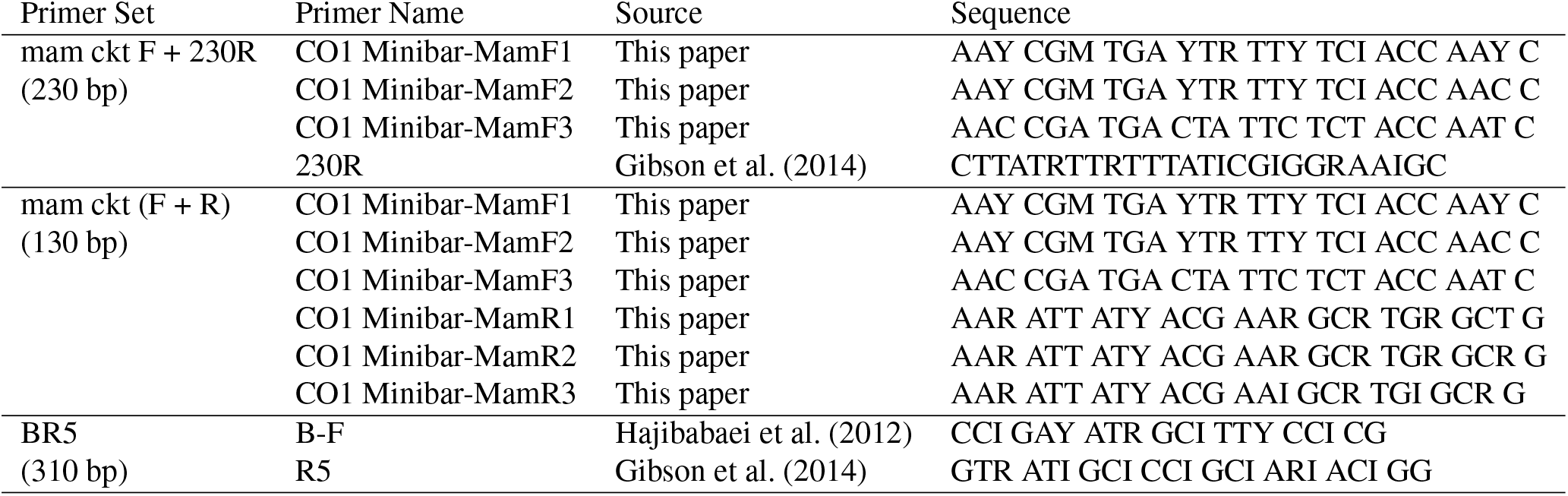

The PCR cycler conditions for each of the cocktails are as follows:

- mam ckt F + 230R: 94°C for 1m, five cycles of (94°C for 30s, 48°C for 40s, 72°C for 1m) and 30 cycles of (94°C for 30s, 50°C for 40s, 72°C for 1m) 72° for 5m, hold at 4°C.
- mam ckt F + R: 94°C for 1m, five cycles of (94°C for 30s, 50°C for 40s, 72°C for 1m) and 35 cycles of (94°C for 30s, 55°C for 40s, 72°C for 1m) 72° for 5m, hold at 4°C
- BR5: 95°C for 5min, 35 cycles of 94°C for 40s, 46°C for 1min, and 72°C for 30s, and a final extension at 72°C for 5min

#### 1.3 Statistical Model for detection by eDNA

We used a hierarchical Bayesian regression with Bernoulli response to model species detections by eDNA metabarcoding across samples (species by sample detections denoted by i subscript). As predictors we included the total visitation per species per sample measured as the numbers of individuals identified at the waterhole edge across all photos (“N waterhole”), the amount of time between sampling and the last instance each species visited a site (“last visit”), using the camera trap data and assuming no observation error. We additionally incuded species average female body mass in kilograms (“mass”) taken from the PanTHERIA mammal trait database (Jones et al., 2009), and sample level environmental predictors including temperature, conductivity, dissolved oxygen, and pH. In addition to these continuous predictors, we included hierarchical predictors for site and species effects, each with adaptively regularizing priors to allow for partial pooling across respective groups. The statistical model follows:

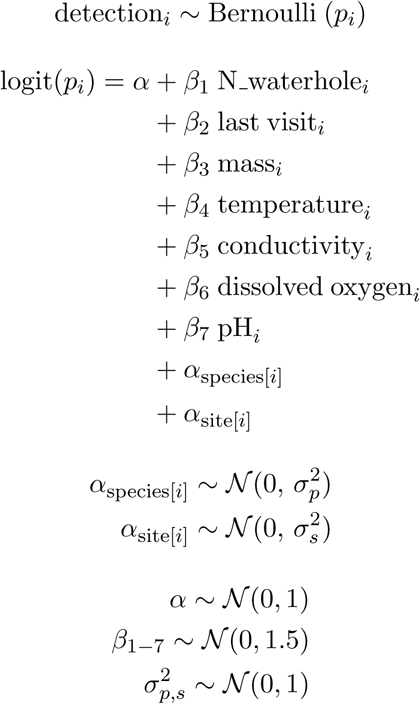

## Supplementary Results

**Table SM2:** Table of species identified by eDNA metabarcoding, according to taxonomic assignment method (i.e. according to the KNP reference library at both the 50 (KNP50) and 80 % minimum bootstrap values, and the Porter Chordata reference library (TP) and the MIDORI reference library (MID) at 80% minimum bootstrap values.)

| Method | Class | Species |
| --- | --- | --- |
| KNP50 | Actinopterygii | *Micropanchax johnstoni* |
| KNP50 | Amphibia | *Tomopterna cryptotis* |
| KNP50 | Aves | *Anhinga rufa* |
| KNP50 | Aves | *Butorides striatus* |
| KNP50 | Aves | *Cinnyris mariquensis* |
| KNP50 | Aves | *Gyps africanus* |
| KNP50 | Aves | *Numida meleagris* |
| KNP50 | Aves | *Sagittarius serpentarius* |
| KNP50 | Aves | *Struthio camelus* |
| KNP50 | Mammalia | *Aepyceros melampus* |
| KNP50 | Mammalia | *Atilax paludinosus* |
| KNP50 | Mammalia | *Ceratotherium simum* |
| KNP50 | Mammalia | *Crocota crocuta* |
| KNP50 | Mammalia | *Elephantulus brachyrhynchus* |
| KNP50 | Mammalia | *Equus quagga* |
| KNP50 | Mammalia | *Gerbilliscus leucogaster* |
| KNP50 | Mammalia | *Giraffa camelopardalis* |
| KNP50 | Mammalia | *Hippopotamus amphibius* |
| KNP50 | Mammalia | *Loxodonta africana* |
| KNP50 | Mammalia | *Neamblysomus julianae* |
| KNP50 | Mammalia | *Panthera leo* |
| KNP50 | Mammalia | *Panthera pardus* |
| KNP50 | Mammalia | *Phacochoerus africanus* |
| KNP50 | Mammalia | *Rhinolophus hildebrandtii* |
| KNP50 | Mammalia | *Sylvicapra grimmia* |
| KNP50 | Mammalia | *Syncerus caffer* |
| KNP50 | Mammalia | *Tragelaphus scriptus* |
| KNP50 | Mammalia | *Tragelaphus strepsiceros* |
| KNP50 | Reptilia | *Prosymna stuhlmannii* |
| KNP50 | Reptilia | *Psammophis mossambicus* |
| KNP80 | Aves | *Gyps africanus* |
| KNP80 | Aves | *Numida meleagris* |
| KNP80 | Mammalia | *Aepyceros melampus* |
| KNP80 | Mammalia | *Ceratotherium simum* |
| KNP80 | Mammalia | *Crocota crocuta* |
| KNP80 | Mammalia | *Equus quagga* |
| KNP80 | Mammalia | *Giraffa camelopardalis* |
| KNP80 | Mammalia | *Hippopotamus amphibius* |
| KNP80 | Mammalia | *Loxodonta africana* |
| KNP80 | Mammalia | *Panthera leo* |
| KNP80 | Mammalia | *Panthera pardus* |
| KNP80 | Mammalia | *Phacochoerus africanus* |
| KNP80 | Mammalia | *Sylvicapra grimmia* |
| KNP80 | Mammalia | *Syncerus caffer* |
| KNP80 | Mammalia | *Tragelaphus scriptus* |
| KNP80 | Mammalia | *Tragelaphus strepsiceros* |
| MID | Actinopterygii | *Zoarces gillii* |
| MID | Amphibia | *Xenopus muelleri* |
| MID | Aves | *Francolinus squamatus* |
| MID | Aves | *Gyps africanus* |
| MID | Aves | *Numida meleagris* |
| MID | Mammalia | *Ceratotherium simum* |
| MID | Mammalia | *Crocota crocuta* |
| MID | Mammalia | *Equus burchellii* |
| MID | Mammalia | *Homo sapiens* |
| MID | Mammalia | *Hystrix cristata* |
| MID | Mammalia | *Loxodonta africana* |
| MID | Mammalia | *Panthera leo* |
| MID | Mammalia | *Panthera pardus* |
| TP | Amphibia | *Xenopus muelleri* |
| TP | Aves | *Francolinus squamatus* |
| TP | Aves | *Gyps africanus* |
| TP | Aves | *Numida meleagris* |
| TP | Mammalia | *Aepyceros melampus* |
| TP | Mammalia | *Equus asinus* |
| TP | Mammalia | *Giraffa camelopardalis* |
| TP | Mammalia | *Homo sapiens* |
| TP | Mammalia | *Hystrix cristata* |
| TP | Mammalia | *Phacochoerus africanus* |
| TP | Mammalia | *Sylvicapra grimmia* |
| TP | Mammalia | *Syncerus caffer* |
| TP | Mammalia | *Tragelaphus scriptus* |
| TP | Mammalia | *Tragelaphus strepsiceros* |

**Figure SM1:**
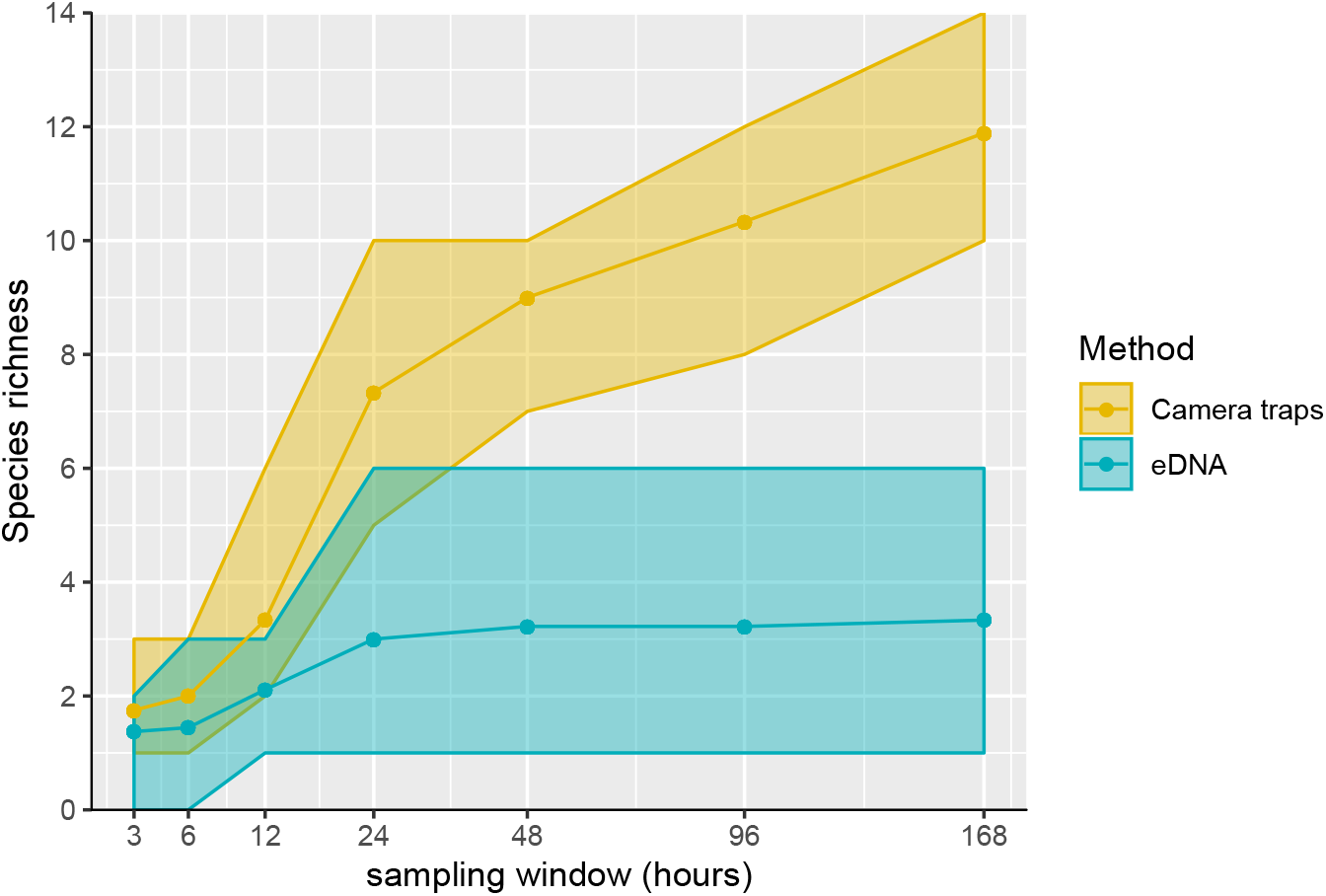
Species richness by camera traps (yellow) and compared with the number of species also detected by eDNA metabarcoding (blue). Richness is calculated per site and across increasing sampling windows starting from the time of first sampling (e.g. 12 hours would represent all animals seen by camera traps within the first 12 hours after sampling a given site, and the number of these same species also detected in the correspnding eDNA sample). As the time between sampling events increases, camera traps detect increasing numbers of species, whereas the number of unique species also detected by eDNA metabarcoding plateaus after roughly 24 hours, indicating that it may be most useful for detecting short-term visitation patterns.

**Table SM3:** Estimated model parameters for eDNA detection model

|  | mean | sd | 2.5% | 50% | 97.5% | effective sample size | Rhat |
| --- | --- | --- | --- | --- | --- | --- | --- |
| log (N waterhole) | 0.11 | 0.17 | -0.22 | 0.10 | 0.46 | 22716.46 | 1.00 |
| sqrt (last visit) | -0.40 | 0.14 | -0.70 | -0.39 | -0.13 | 26697.80 | 1.00 |
| log (mass) | 0.30 | 0.21 | -0.07 | 0.29 | 0.73 | 19399.64 | 1.00 |
| Temperature | 0.01 | 0.29 | -0.53 | 0.01 | 0.59 | 23519.40 | 1.00 |
| Conductivity | 0.55 | 0.38 | -0.24 | 0.55 | 1.27 | 17024.15 | 1.00 |
| Dissolved Oxygen | 0.06 | 0.18 | -0.27 | 0.06 | 0.44 | 15372.11 | 1.00 |
| pH | 0.12 | 0.87 | -1.58 | 0.13 | 1.83 | 22765.78 | 1.00 |
| sigma species | 0.61 | 0.41 | 0.03 | 0.55 | 1.54 | 8048.31 | 1.00 |
| sigma site | 0.54 | 0.45 | 0.02 | 0.43 | 1.67 | 8818.97 | 1.00 |

To quantify variation in taxonomic assignment across methods, we partitioned Sorensen’s beta diversity across the species lists generated by each approach and find that the majority of differences across approaches is due to turnover rather than nestedness (Fig. SM2). This means that across the reference libraries, different species are being identified rather than each method identifying smaller subsets of the same set of species.

**Figure SM2:**
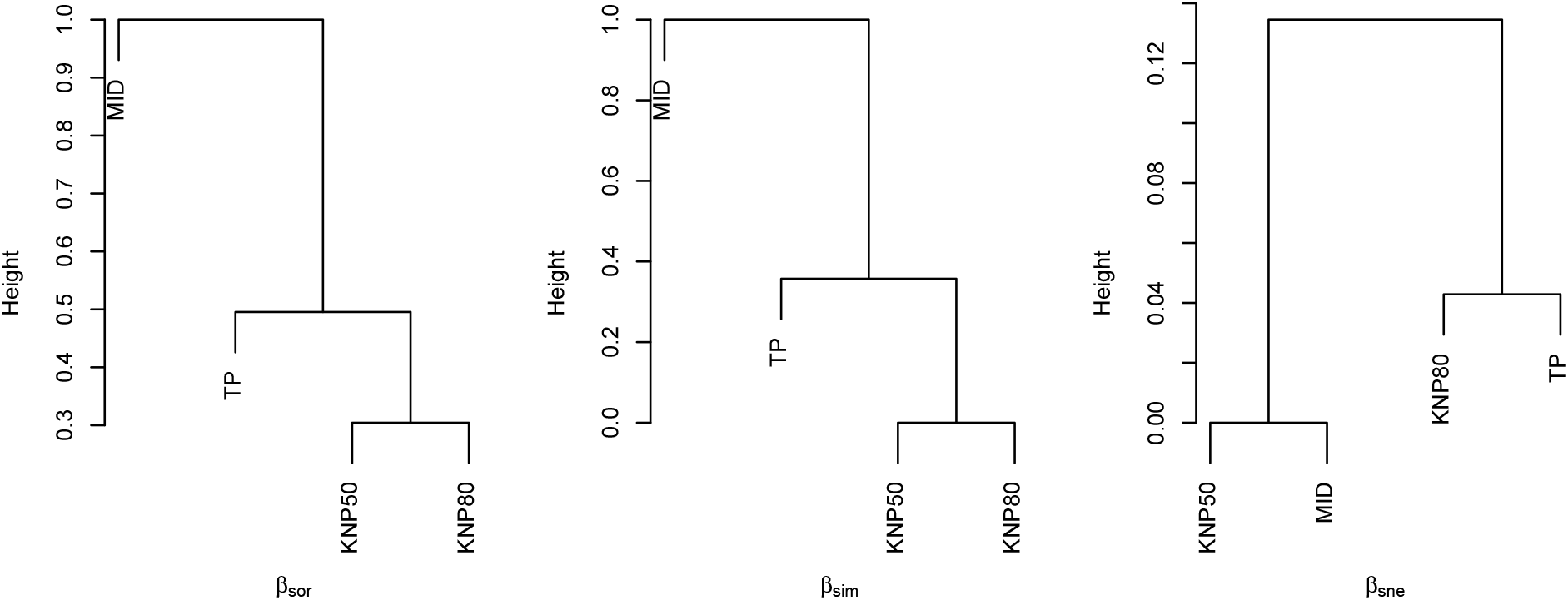
Similarity in species identified via eDNA metabarcoding as a function of reference library approach. Differences between sets of species identified with each method are analyzed by hierarchical clustering using Sorensen’s beta diversity (*β*_*sor*_), which breaks down into a turnover component (Simpson’s diversity (*β*_*sim*_), and a nestedness (*β*_*sne*_) component. This approach shows that the two KNP reference library approaches (KNP50 and KNP80) identify similar species, which more closely resembles the results of the Porter Chordata reference library (TP) compared to the MIDORI reference library (MID). The other two metrics indicate that this similarity is largely attributed turnover rather than nestedness, indicating that the KNP reference library is identifying different species, rather than a subset of species identified by the other approaches. Analyses conducted using the “betapart” R package (Baselga and Orme, 2012).

**Figure SM3:**
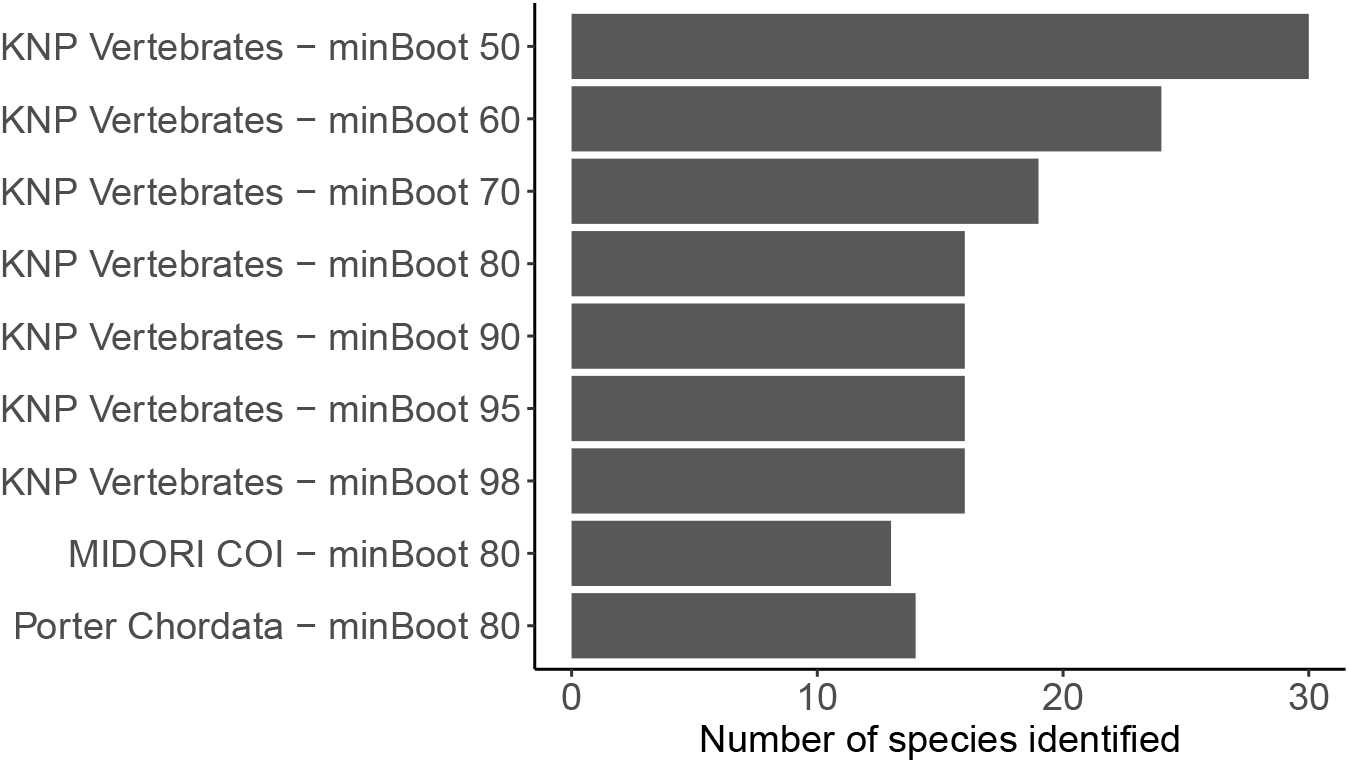
Total vertebrate species richness identified by eDNA metabarcoding across all samples according to taxonomic reference library (Kruger Reference Library (KNP), MIDORI COI (MID), and Porter Chordata (TP), and the minimum bootstrap (minBoot) value used when applying the RDP classifier via dada2.

**Table SM4:** Per species rates of eDNA (KNP Vertebrate Reference Library with RDP classifier at 80% minimum bootstrap) and camera detections by sample. * denotes species which did not have representative sequences in the KNP vertebrate reference library. Across all species that had reference sequences, 25% of species by sample combinations identified by camera trapping were recovered by eDNA.

| Species | eDNA detections | Camera detections | Rate |
| --- | --- | --- | --- |
| *Crocota crocuta* | 5 | 8 | 0.62 |
| *Ceratotherium simum* | 3 | 6 | 0.50 |
| *Panthera pardus* | 1 | 2 | 0.50 |
| *Aepyceros melampus* | 4 | 9 | 0.44 |
| *Loxodonta africana* | 4 | 9 | 0.44 |
| *Hippopotamus amphibius* | 2 | 5 | 0.40 |
| *Equus quagga* | 2 | 7 | 0.29 |
| *Syncerus caffer* | 1 | 5 | 0.20 |
| *Panthera leo* | 1 | 6 | 0.17 |
| *Tragelaphus strepsiceros* | 1 | 6 | 0.17 |
| *Giraffa camelopardalis* | 1 | 7 | 0.14 |
| *Phacochoerus africanus* | 1 | 9 | 0.11 |
| *Connochaetes taurinus* | 0 | 7 | 0.00 |
| *Canis mesomelas* | 0 | 5 | 0.00 |
| *Chlorocebus pygerythrus* | 0 | 4 | 0.00 |
| *Civettictis civetta* | 0 | 4 | 0.00 |
| *Papio ursinus* | 0 | 4 | 0.00 |
| *Kobus ellipsiprymnus* | 0 | 1 | 0.00 |
| * *Mungos mungo* | 0 | 1 | 0.00 |
| * *Paraxerus cepapi* | 0 | 1 | 0.00 |
| *Tragelaphus scriptus* | 0 | 1 | 0.00 |

**Table SM5:** Per species rates of eDNA and camera detections by sample, subset to camera trap photographs taken within 12 hours prior to sampling for eDNA. Across all species 50% of species by sample combinations identified by camera trapping were recovered by eDNA.

| Species | eDNA detections | Camera detections | Rate |
| --- | --- | --- | --- |
| *Hippopotamus amphibius* | 1 | 1 | 1.00 |
| *Syncerus caffer* | 1 | 1 | 1.00 |
| *Crocuta crocuta* | 4 | 6 | 0.67 |
| *Ceratotherium simum* | 3 | 5 | 0.60 |
| *Aepyceros melampus* | 2 | 4 | 0.50 |
| *Equus quagga* | 2 | 4 | 0.50 |
| *Loxodonta africana* | 2 | 4 | 0.50 |
| *Canis mesomelas* | 0 | 3 | 0.00 |
| *Civettictis civetta* | 0 | 1 | 0.00 |
| *Giraffa camelopardalis* | 0 | 1 | 0.00 |

**Figure SM4:**
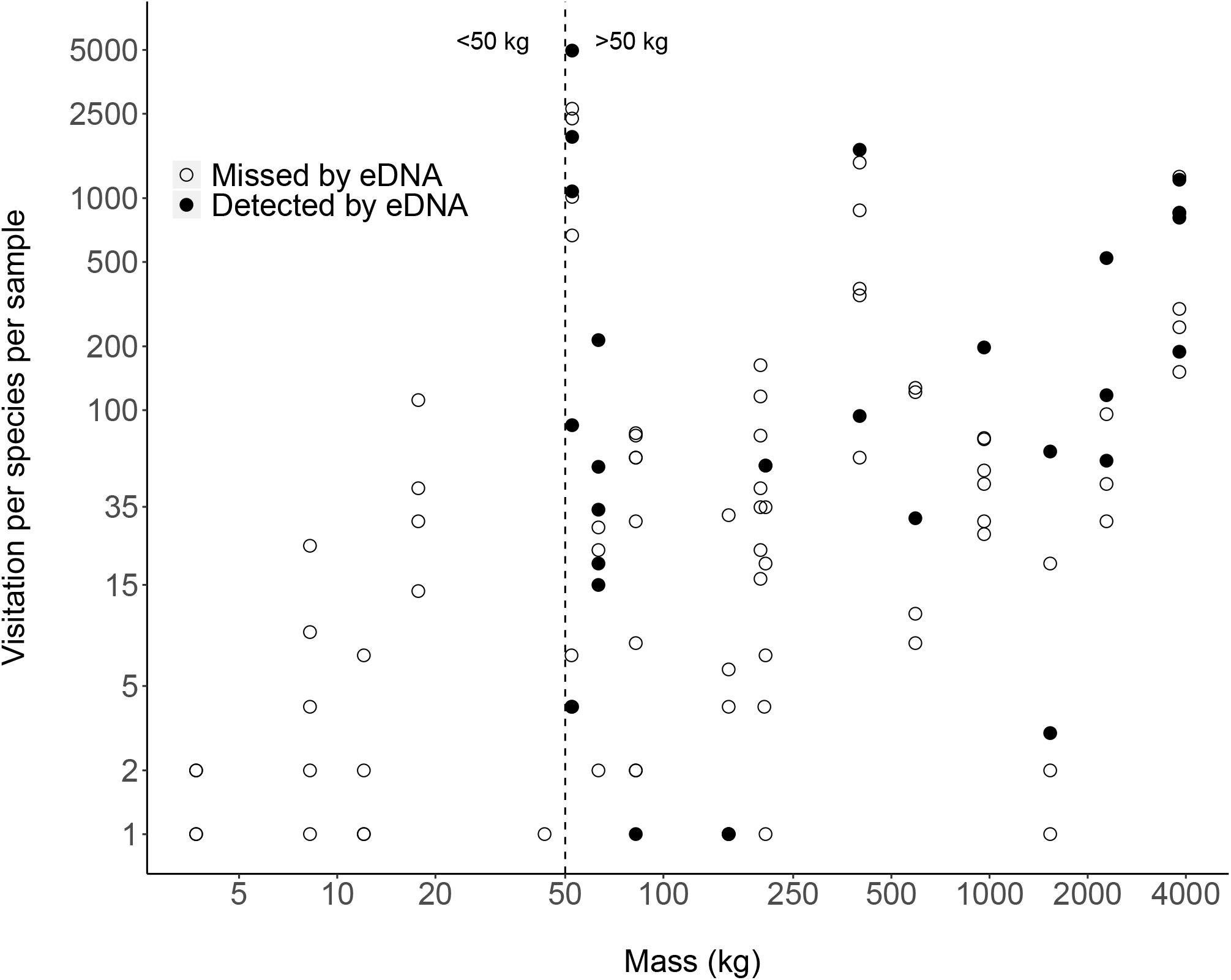
Raw data plot showing the relationship between species mass (average adult female mass in kilograms) and the total number of animals of that species obverved via camera trapping. Filled points indicate samples which have positive eDNA detection for the given species.

**Figure SM5:**
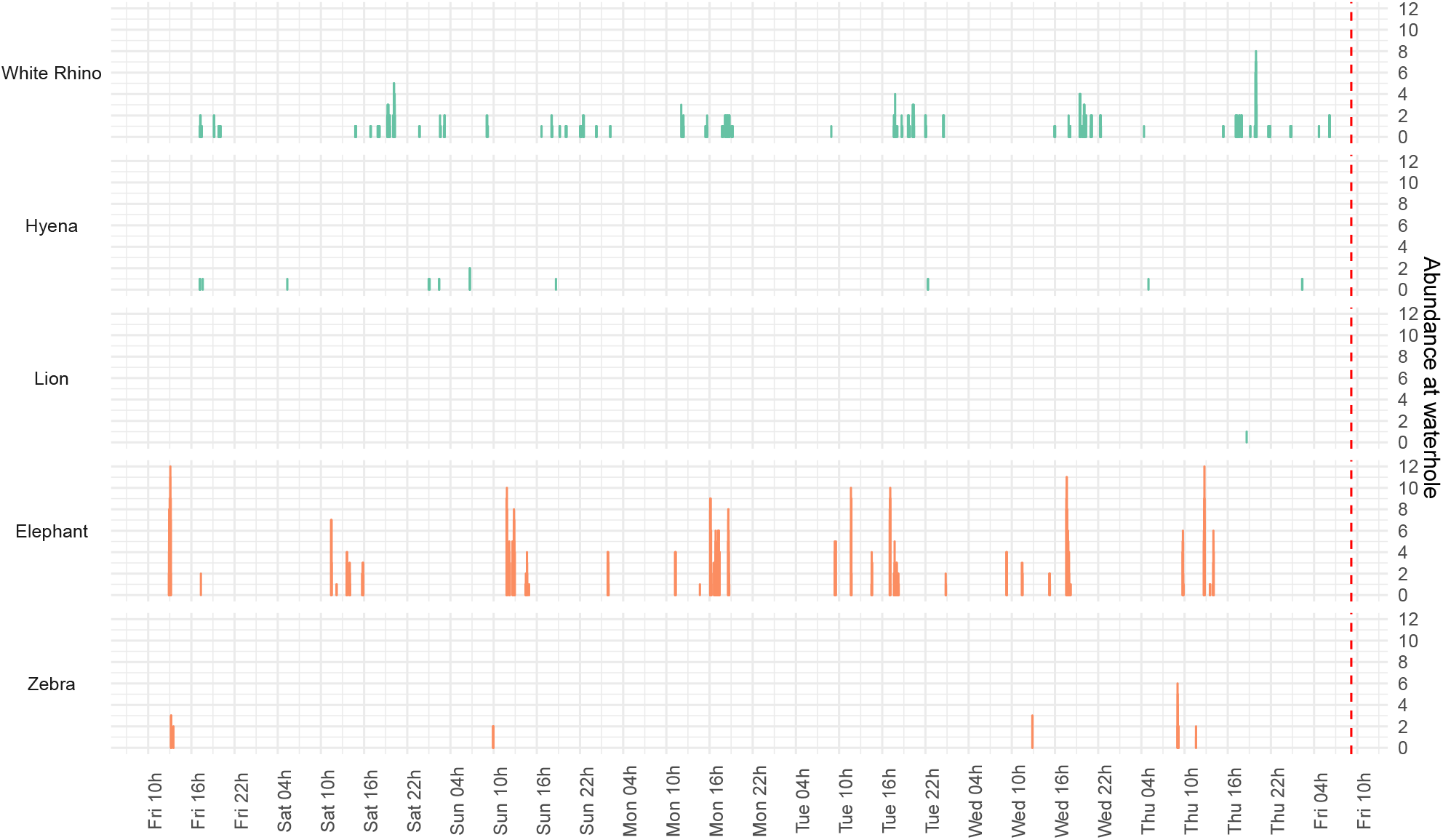
Example of visitation timeseries for five species based one week of camera trapping. Data from a single site is used for demonstration. The red dashed line indicates the time water was sampled. Orange lines indicate abundances of species that were seen in the camera traps but not detected by eDNA, while turquiose lines represent species detected by both camera trapping and eDNA, as in Fig. 3

**Figure SM6:**
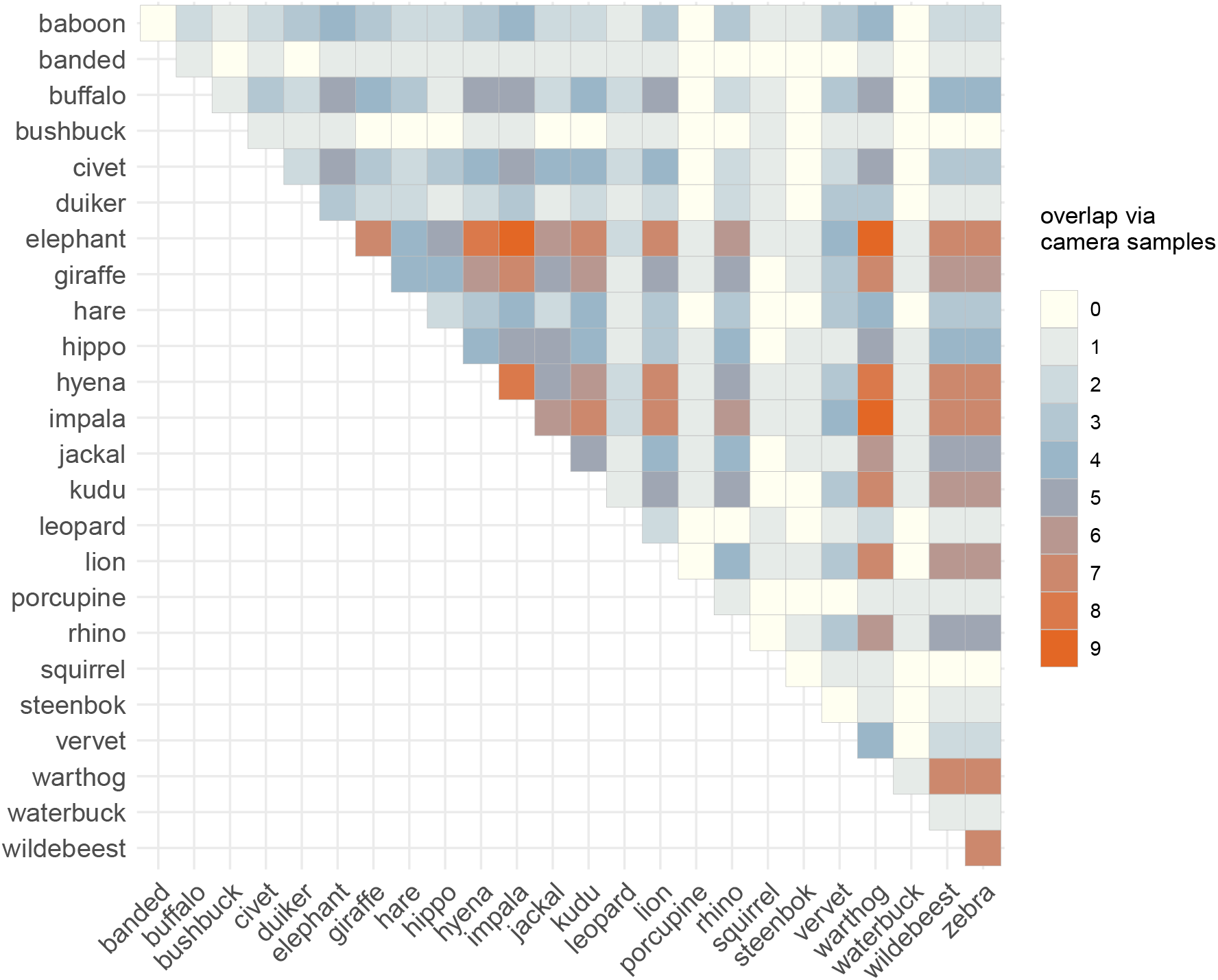
Heatmap of camera-trap observed overlap among pairs of mammals. Overlap among species pairs is quantified by the number of weekly samples for which both species were documented by camera traps. For example, impalas and elephants were seen at the same sites in 9 weekly samples whereas baboons and banded mongeese were never seen at any of the same sites.

**Figure SM7:**
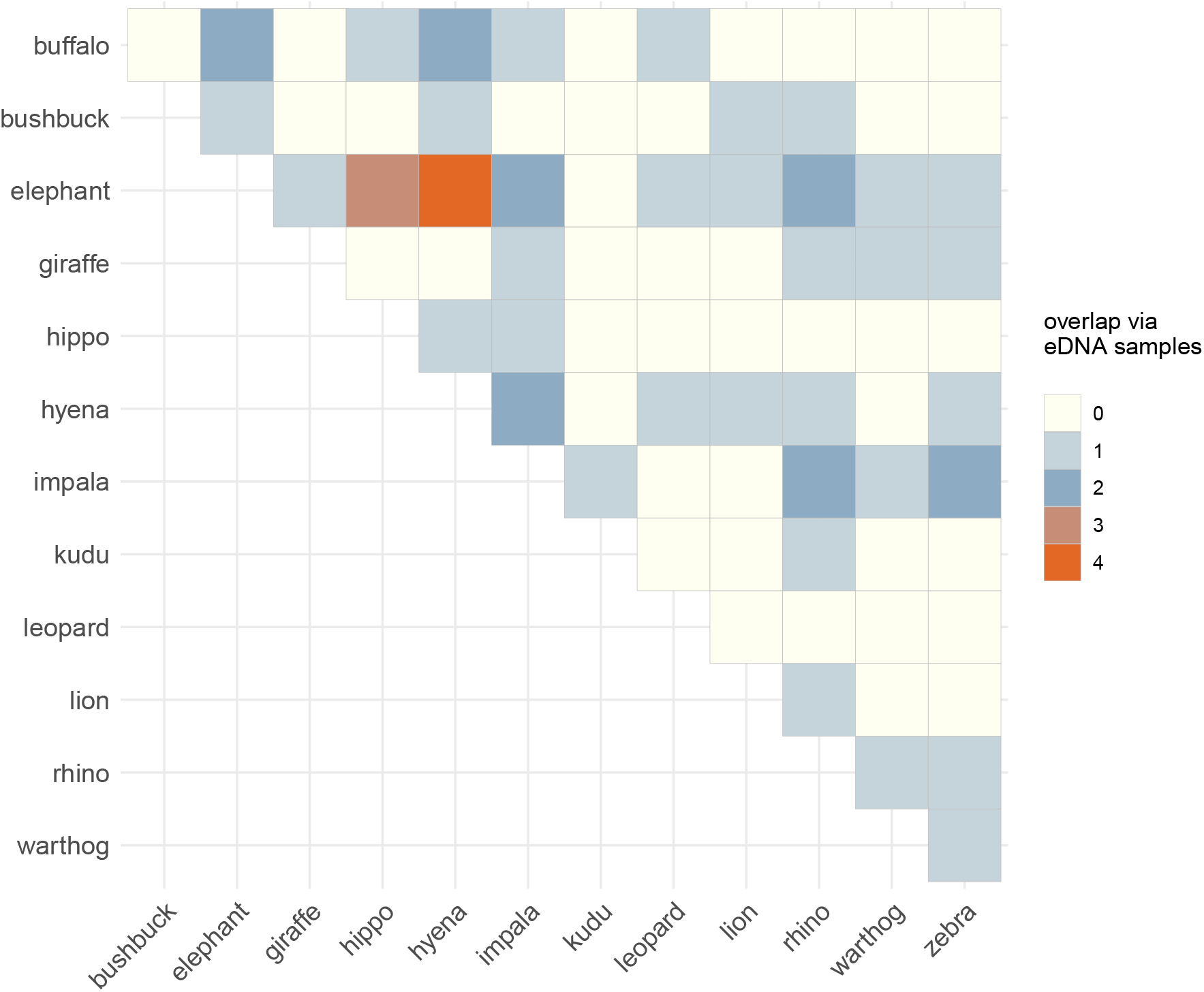
Heatmap of eDNA inferred pairwise overlap among species present at waterholes. Overlap among species pairs is quantified by the number of eDNA samples in which metabarcoding simultaneously detected both species. For example, eDNA from elephants and impala were detected in 4 weekly samples, whereas eDNA from baboons and banded mongeese were never detected in any of the same samples.

